# Transposon Screen of Surface Accessibility in *S. aureus*

**DOI:** 10.1101/2021.09.03.458708

**Authors:** Noel J. Ferraro, Marcos M. Pires

## Abstract

Bacterial cell walls represent one of the most prominent targets of antibacterial agents. These agents include natural products (e.g., vancomycin) and proteins stemming from the innate immune system (e.g., peptidoglycan-recognition proteins and lysostaphin). Among bacterial pathogens that infect humans, *Staphylococcus aureus* (*S. aureus*) continues to impose a tremendous healthcare burden across the globe. *S. aureus* has evolved countermeasures that can directly restrict the accessibility of innate immune proteins, effectively protecting itself from threats that target key cell well components. We recently described a novel assay that directly reports on the accessibility of molecules to the peptidoglycan layer within the bacterial cell wall of *S. aureus*. The assay relies on site-specific chemical remodeling of the peptidoglycan with a biorthogonal handle. Here, we disclose the application of our assay to a screen of a nonredundant transposon mutant library for susceptibility of the peptidoglycan layer with the goal of identifying genes that contribute to the control of cell surface accessibility. We discovered several genes that resulted in higher accessibility levels to the peptidoglycan layer and showed that these genes modulate sensitivity to lysostaphin. These results indicate that this assay platform can be leveraged to gain further insight into the biology of bacterial cell surfaces.

**Table of Contents Figure:** 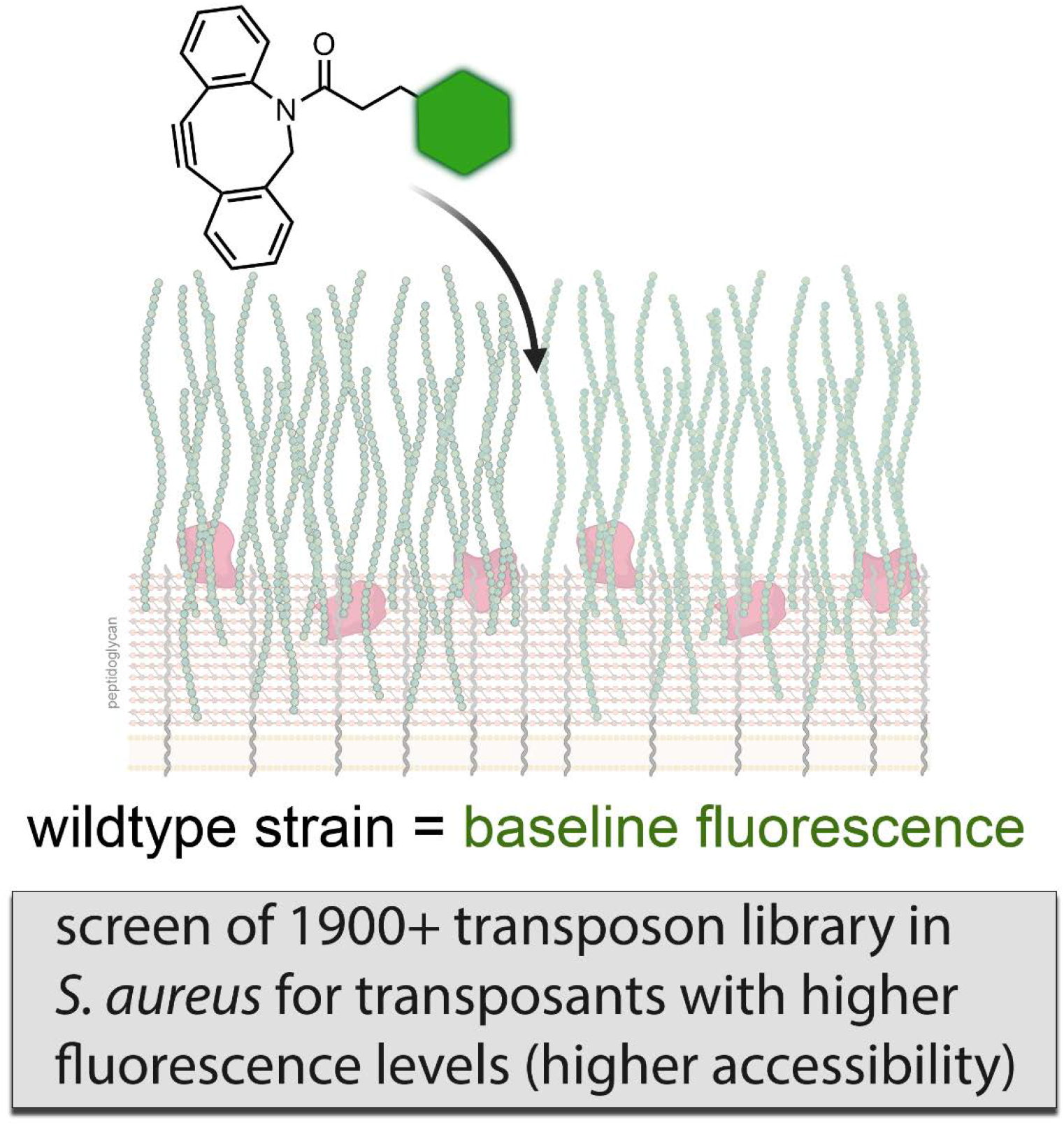

## Introduction

Bacterial resistance to antibiotics has become an imminent threat to global health and must be met with an antibiotic pipeline revitalization or, *in lieu* of that, alternative methods to combat bacterial infections. Intrinsic resistance is a multifactorial phenomenon but sometimes it can be mediated by a simple lack of accessibility to the bacterial targets.^*1–3*^ Recent efforts have led to the discovery of molecules that can potentiate antibiotics by improving permeation to and across the bacterial cell surface. For example, Gram-negative pathogens can be sensitized by co-treatment with polymyxins (and similar outer membrane destabilizers), which improve permeation of antimicrobials through the bacterial cell wall.^*4–8*^ Likewise, positively charged Branched PolyEthylenImine (BPEI) has been shown to potentiate β-lactam antibiotics against the Gram-positive pathogen methicillin resistant *Staphylococcus aureus* (MRSA).^*9, 10*^ It was proposed that BPEI neutralization of negatively charged polymers on the bacterial cell surface improved permeability.^*11*^ These examples demonstrate that improved permeation to the essential cell wall components can be a powerful modality of reducing intrinsic resistance to small molecule antibacterials and immune proteins.^*12*^

The cell walls of bacteria are complex in structure and composition. For the Gram-positive bacterium, *Staphylococcus aureus* (*S. aureus*), the cell wall is composed of a thick peptidoglycan (PG) scaffold that is shielded by proteins and other biomacromolecules (**Figure 1**).^*13*^ The proteins are covalently anchored onto the PG *via* the transpeptidase, sortase A.^*14, 15*^ The cell surface is further functionalized by wall teichoic acids (WTAs), which are anionic glycopolymers that are covalently anchored onto the PG, and lipoteichoic acids (membrane anchored).^*16*^ The restricted accessibility afforded by teichoic acids and other surface-bound macromolecules has been implicated in the virulence of Staphylococci. In *Drosophila*, it was recently discovered that the accessibility of peptidoglycan recognition proteins (PGRPs) to PG plays a determinant role in the host immunity to infection.^*17*^ Removal of WTAs resulted in considerably increased accessibility of PGRPs to the PG and predisposed *S. aureus* to PGRP-mediated immunity. Additionally, it has been found that WTA-deficient *S. aureus* fails to colonize the nasal cavities of rats.^*18*^ More recently, it was established that *Staphylococcus epidermidis* shift from a commensal to pathogen lifestyle upon expression of *S. aureus*-like WTAs.^*19*^ Aside from modulating immunity, PG accessibility can potentially impact drug discovery based on the fact that a number of FDA approved antibiotics work by inhibiting steps in the bacterial PG biosynthesis pathway.^*20, 21*^ The essential nature of accessibility to the PG scaffold by immune proteins and antibiotics alike means that there is a need to better understand the genes that control surface accessibility.

**Figure 1.**
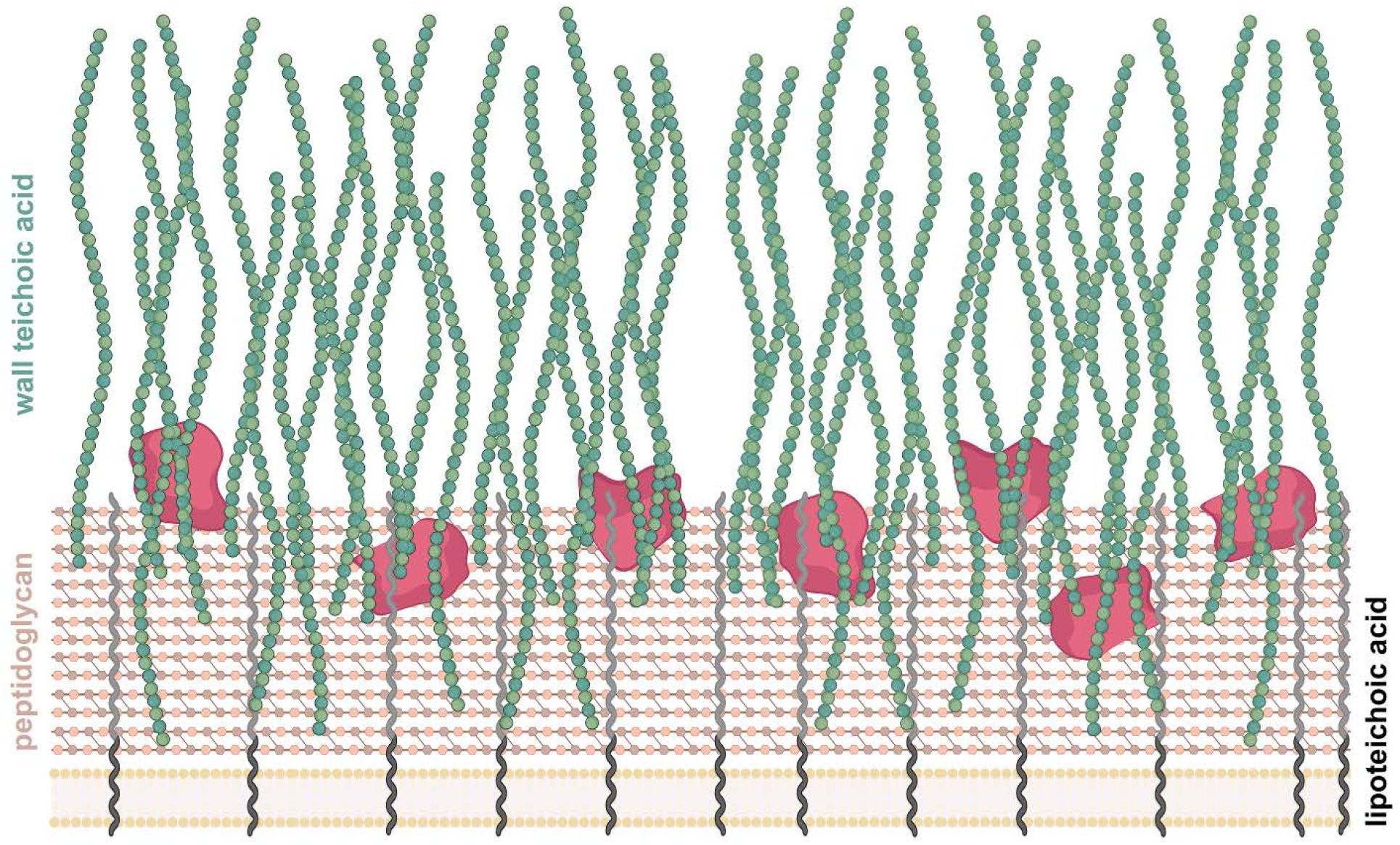
Schematic representation of the surface of Gram-positive bacteria. The PG scaffold is heavily decorated with a number of biomacromolecules, including WTAs, LTAs, and proteins. These features can reduce the permeation of extracellular proteins and molecules to the PG scaffold, which can modulate host immunity and the potency of antibacterial agents.

Among Gram-positive pathogens, *S. aureus* has proven to be particularly challenging to treat; it is a formidable foe, as it is well suited to evade attacks by the host immune system^*22*^ and it is a pathogen that can readily become resistant to standard of care therapies.^*23*^ The difficulty in finding new efficacious antibiotics against *S. aureus* highlights the need to explore less conventional therapeutic approaches such as antibiotic adjuvants or immunotherapies.^*24–29*^ For example, adjuvants can potentiate antibiotics by improving their accessibility to their cognate molecular targets. Likewise, anti-infective immunotherapeutics (e.g., antibody recruiting agents developed by our lab^*24–27, 29*^) work by targeting specific macromolecules on bacterial cell surfaces. Despite the pivotal role of surface accessibility in bacterial pathogenesis, to date there has not been a systematic analysis of the genes that control penetration of biomacromolecules across cell surfaces of *S. aureus*; information that can directly impact current therapies and the development of alternative treatments. To address this, we performed a screen against a non-redundant transposon mutant library using a robust assay that reports on surface accessibility in *S. aureus*.

## Results and Discussion

We set out to comprehensively screen nonessential genes in *S. aureus* for their ability to alter accessibility to the PG scaffold. The basis of the screening assay is site-specific incorporation of a biorthogonal handle *via* metabolic remodeling of bacterial PG.^*11*^ More specifically, an unnatural amino acid (**D-LysAz**) is supplemented in the media, inoculated with *S. aureus*, and cultured overnight (**Figure 2A**). During PG biosynthesis and assembly, transpeptidases catalyze the replacement of the 5^th^ position D-alanine on the stem peptide within the bacterial PG scaffold for **D-LysAz**, thus leading to the covalent installation of the azido-handle (**Figure 2B**). With the azido group installed through the entire PG scaffold, treatment of cells with a complementary DiBenzoCycloOctyne (DBCO) handle conjugated to fluorescein (**DBCOfl**) leads to the specific tagging of the bacterial PG with fluorescent moieties (**Figure 2C**).^*30, 31*^ We previously showed that cellular fluorescence levels reflect the ability of molecules to reach the PG scaffold and consequently result in a covalent tag. We hypothesized that this reporter assay could be leveraged to identify genes that modulate PG accessibility in *S. aureus*, thus revealing genes that have implications in immunity and drug discovery. By screening a non-redundant library of transposon mutants, we reasoned that any mutants exhibiting increased cellular fluorescence levels could identify potential genetic alterations that result in higher surface accessibility.

**Figure 2.**
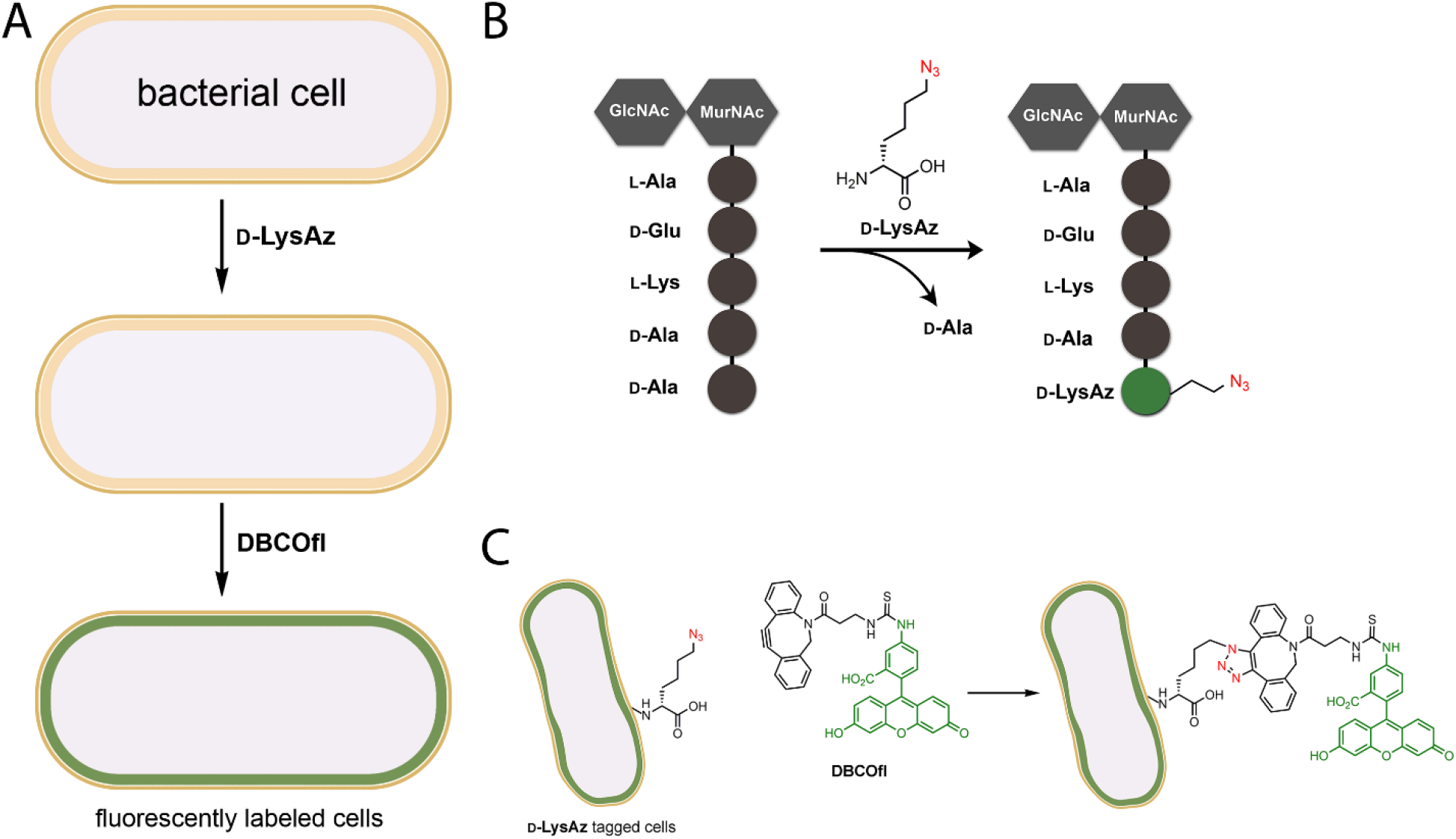
(**A**) Live cell labeling with **D-LysAz** and **DBCOfl**. Cells are analyzed by the flow cytometer to quantify modification with **DBCOfl**. (**B**) The unnatural amino acid (**D-LysAz**) is enzymatically installed in place of the terminal D-alanine on the stem peptide within the bacterial PG during cellular growth. (**C**) Chemical structures depicting the reaction between the azido moiety and DBCO.

First, we demonstrated that the assay preformed as expected in wildtype (WT) *S. aureus* cells by supplementing the PG label (**D-LysAz**) during overnight culture. Subsequent treatment with **DBCOfl** resulted in cellular fluorescence levels that were ~6-fold and ~8-fold higher than no amino acid and the **L-LysAz**, respectively (**Figure 3A**). We^*32, 33*^, and others^*34–37*^, had previously demonstrated that the enantiomeric **L-LysAz** does not incorporate into the growing PG scaffold of bacteria. These results confirm that cellular fluorescence increases in a manner that is dependent on PG remodeling with azido handles. To demonstrate the effect of surface accessibility, a similar labeling protocol was performed with *tarO*-deleted *S. aureus*, which results in WTA-free S. aureus cells.^*16, 38, 39*^ Fluorescence levels of *ΔtarO S. aureus* labeled with **D-LysAz** were ~2-fold higher than WT cells (**Figure 3A**). These results are consistent with previous findings showing that WTAs control the accessibility of immunoproteins to the PG.^*17, 40, 41*^

**Figure 3.**
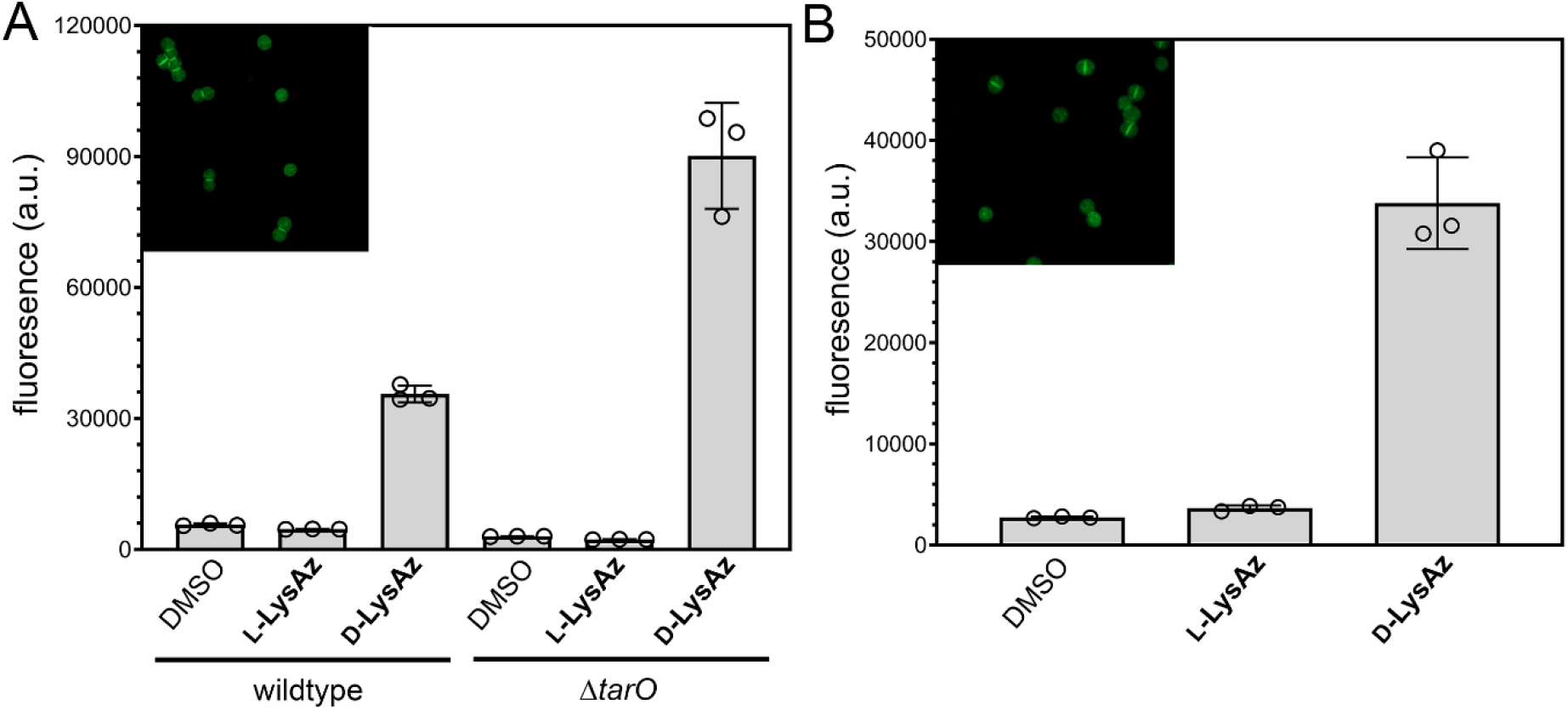
(**A**) Flow cytometry analysis of WT *S. aureus* (ATCC 25923) or *S. aureus* (*ΔtarO*) treated overnight with DMSO, 1 mM of **L-LysAz**, or 1 mM of **D-LysAz** followed by a treatment with 25 μM of **DBCO-FI**. *Inset*, confocal microscopy of the WT cells tagged with and treated with **D-LysAz** followed by a treatment with 25 μM of **DBCOfl**. (**B**) Sacculi isolated from WT *S. aureus* (ATCC 25923). WT cells were incubated with 1 mM of **D-LysAz**, 1 mM of **L-LysAz**, or DMSO alone overnight. Next, the cells were treated with 25 μM of **DBCO-FI** and subjected to a sacculi isolation protocol. The resulting sacculi were analyzed by flow cytometry. *Inset*, confocal microscopy of the sacculi from WT cells that were treated with **D-LysAz** followed by a treatment with 25 μM of **DBCOfl**. Data are represented as mean +/− SD (n = 3).

We then performed an additional set of experiments to confirm that the fluorescent reporter handle reached the PG scaffold. Our research lab recently described an assay (SaccuFlow) that combines the quantification of flow cytometry with the analysis of bacterial sacculi.^*11*^ Similar to the whole cell assay described in **Figure 2**, *S. aureus* cells were treated with **D-LysAz** followed by **DBCOfl**. Instead of analyzing the whole cell, the sacculus was isolated using a standard isolation procedure which includes boiling of cells with SDS (to solubilize biomacromolecules) and treatment with trypsin (to remove proteins anchored on the PG scaffold). Following these steps, the isolated sacculi are analyzed on the flow cytometer. As expected, fluorescence levels of sacculi from cells treated with a combination of **D-LysAz** and **DBCOfl** were ~12-fold and ~9-fold higher than sacculi from cells treated with DMSO or **L-LysAz**, respectively (**Figure 3B**). Confocal microscopy confirmed that the whole cells and the isolated sacculi retained the expected size and shape of the fluorescently labeled cells and biomacromolecule, respectively (insets of **Figure 3**). To confirm the metabolic labeling step did not alter cellular viability, a growth curve analysis was performed by monitoring the optical density at 600 nm (**Figure S1**). Consistent with our prior results using D-amino acid labeling at the concentrations used in our assay, cellular viability was not impacted. Together, these results show that the combination of PG labeling with **D-LysAz** and treatment with the **DBCOfl** in the media results in the fluorescence tagging of bacterial PG in a manner that reports on cell surface accessibility.

We next shifted our efforts to demonstrate the utility of the surface accessibility assay by screening a library of *S. aureus* transposon mutants to identify genes that could play important roles in controlling permeation to the *S. aureus* cell surface. To this end, a transposon (*Tn*) insertion mutant library was used as the platform of the screen, which contained 1952 individual strains each with a single insertion within a nonessential gene in *S. aureus* USA300. A major advantage of using the transposon mutant library is the potential to identify genes that were not previously purported to have biological roles related to accessibility, thus revealing new targets for the development of adjuvant agents. Before screening the entire library, the assay was benchmarked in a 384-well format and it was found that the assay could be readily miniaturized (**Figure S2**). Next, the entire library was screened using the combination of **D-LysAz** and **DBCOfl** (**Figure 4A**). While most transposon mutants did not significantly display altered labeling levels, several transposants exhibited elevated fluorescence levels (defined as 33%+ above the average and denoted by the green line). These results suggest that those genes may potentially play a role in surface accessibility.

**Figure 4.**
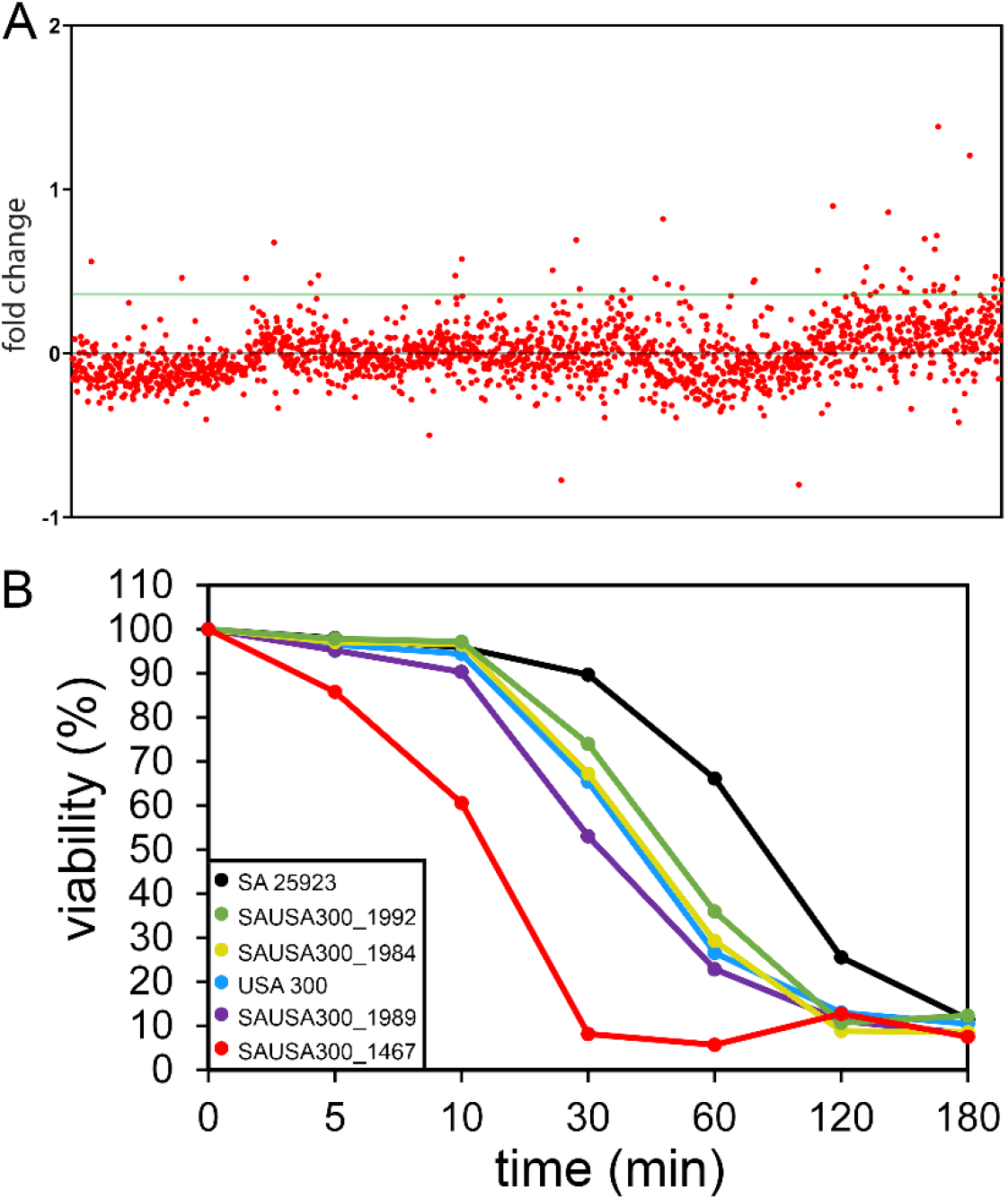
**(A)** Flow cytometry analysis of transposon library in *S. aureus* (ATCC 25923) treated overnight with 1 mM of **D-LysAz**, followed by a treatment with 25 μM of **DBCOfl**. Fluorescence levels were normalized to the average levels across all transposon mutants. Fold change refers to the ratio of fluorescence of the strain relative to the library average. Black line represents the zero point (average) and the green line represents +0.33-fold change above the average, which we designated as potential hits. **(B)** Percent viability of designated transposon mutants and wild-type strains of *S. aureus* when challenged with 5 μg/mL of lysostaphin. Measurement of cellular viability was performed by analyzing the optical density at 600 nm and at each time point the optical density value was compared to the initial optical density reading.

The 24 transposants with the highest increase in cellular fluorescence were re-assayed in triplicate and transposon mutants that consistently labeled at a higher level than wild-type were selected (**Figure S3**). These results showed that four transposants were confirmed to play a determinant role in the accessibility of **DBCOfI** to the cell surface, namely SAUSA300_1989, SAUSA300_1984, SAUSA300_1992, and SAUSA300_1467. SAUSA300_1989 and SAUSA300_1992 have been annotated as *agrB* and agrA, respectively.^*42*^ Their protein products play a central role in processing autoinducing peptides in *S. aureus*. Relatedly, the protein product of SAUSA300_1984 (*mroQ*) acts within the Agr pathway.^*43*^ Moreover, the protein product of SAUSA300_1467 (*lpdA*) has been shown to have a role in branched-chain fatty acid biosynthesis.^*44*^ The selected transposants were also tested for labeling with a second fluorescent probe (**DBCO-AF488**) in order to determine the role of the fluorophore in PG labeling (**Figure S4**). The labeling pattern of the **DBCO-AF488** mirrored that of **DBCOfI** with the mutants labeling at a higher level than WT. Similarly, we evaluated the impact of a spacer length between the DBCO handle and the fluorescein moiety across the transposants. A polyethylene glycol spacer was inserted between DBCO and fluorescein (**DBCOpegfl**) to test how a larger molecule would compare to the smaller **DBCOfl**. It was observed that increasing the size of the probe resulted in similar profile to that of **DBCOfI**, which indicates that the effect of increased fluorescence in the transposants as compared to WT is relevant for different sized probes (**Figure S5**).

Finally, the identified hits were tested for their susceptibility to a larger molecular weight protein, lysostaphin, which needs to reach the PG layer to impart its antimicrobial activity (**Figure 4B**). We reasoned that genes that impact surface accessibility may be able to modulate the activity of lysostaphin by increasing its access to its target biomacromolecule, the bacterial PG. Lysostaphin is a bacteriocin that cleaves the pentaglycine cross-bridges found in the cell walls of *S. aureus*, leading to bacterial lysis.^*45*^ Challenge with lysostaphin showed that disruption to those genes resulted in altered sensitivity, which suggests the possibility that surface accessibility is playing a role lysostaphin activity. Together, this pilot screen confirmed the ability of the assay to be miniaturized to formats that are compatible with high-throughput screening and it reveals potential genes that may control surface accessibility of *S. aureus*.

## Conclusion

In conclusion, we have shown the application of our novel assay that reports on accessibility to the PG in identifying genes that potentially control permeation to the bacterial cell surface in *S. aureus*. Using site-selective incorporation of a biorthogonal handle and a corresponding reactive fluorescent probe, we were able to screen a transposon mutant library in effort to discern any genetic variations that led to an increase in fluorescence labeling thus signaling a possible increase in permeability. The screen revealed four transposants that consistently labeled at a higher level when compared to WT cells and some of these identified hits showed increased susceptibility to the bacteriocin, lysostaphin. The protein products of these transposants can potentially be considered new targets to develop potentiators for antibacterial therapies. Overall, we have demonstrated the use of our PG accessibility assay in a high-throughput screen and identified genes that may control access to the cell wall surface in *S. aureus*.

