## Supplemental Figures and Methods for "Transposon Screen of Surface Accessibility in *S. aureus*"

**Supporting Figures**

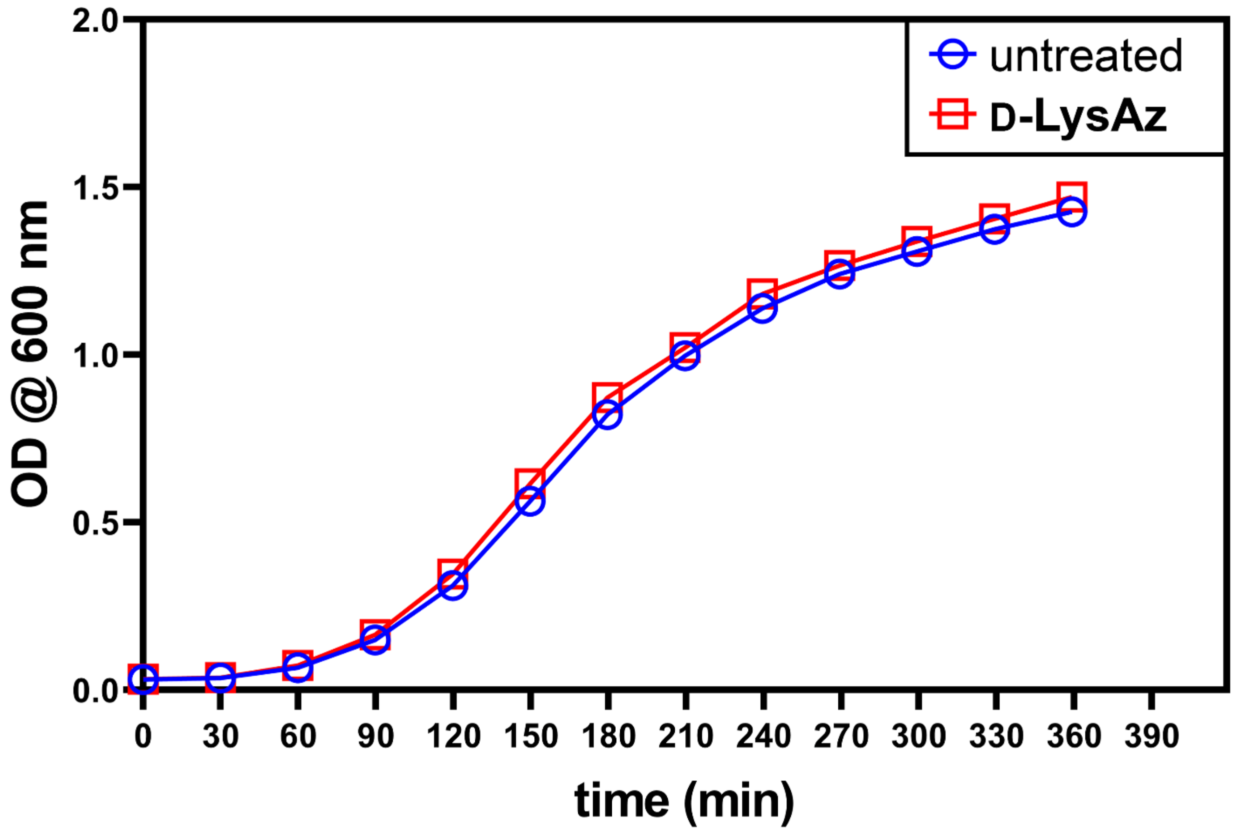


**Figure S1.** Cellular viability of *S. aureus* with treatment of **D-LysAz**. A culture of S. aureus

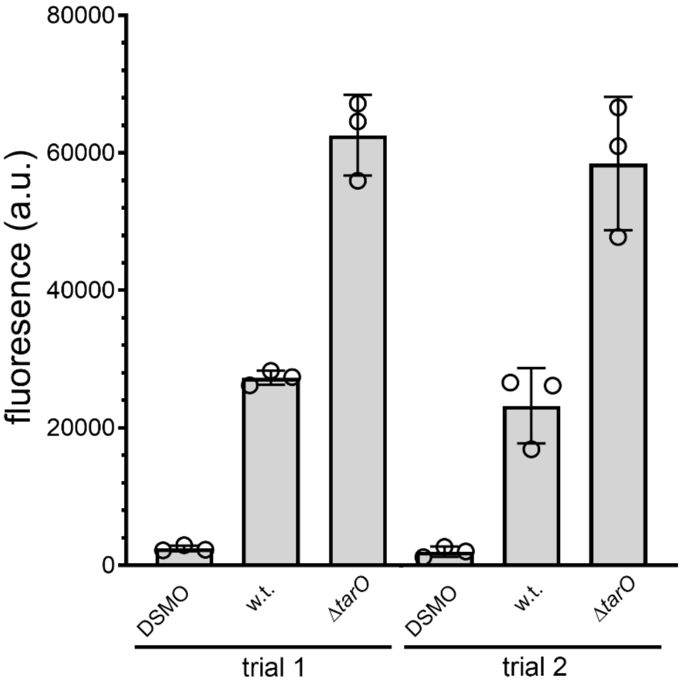


**Figure S2.** Flow cytometry analysis of WT *S. aureus* (ATCC 25923) and *ΔtarO* grown in a 384-well plate, treated overnight with 1 mM of **D-LysAz**, followed by a treatment with 25 μM of **DBCO-Fl**. Cells were analyzed in the 384-well plates using a CytoFlex flow cytometer. Two independent trials were performed to demonstrate the stability of the assay. Data are represented as mean +/- SD (n = 3).

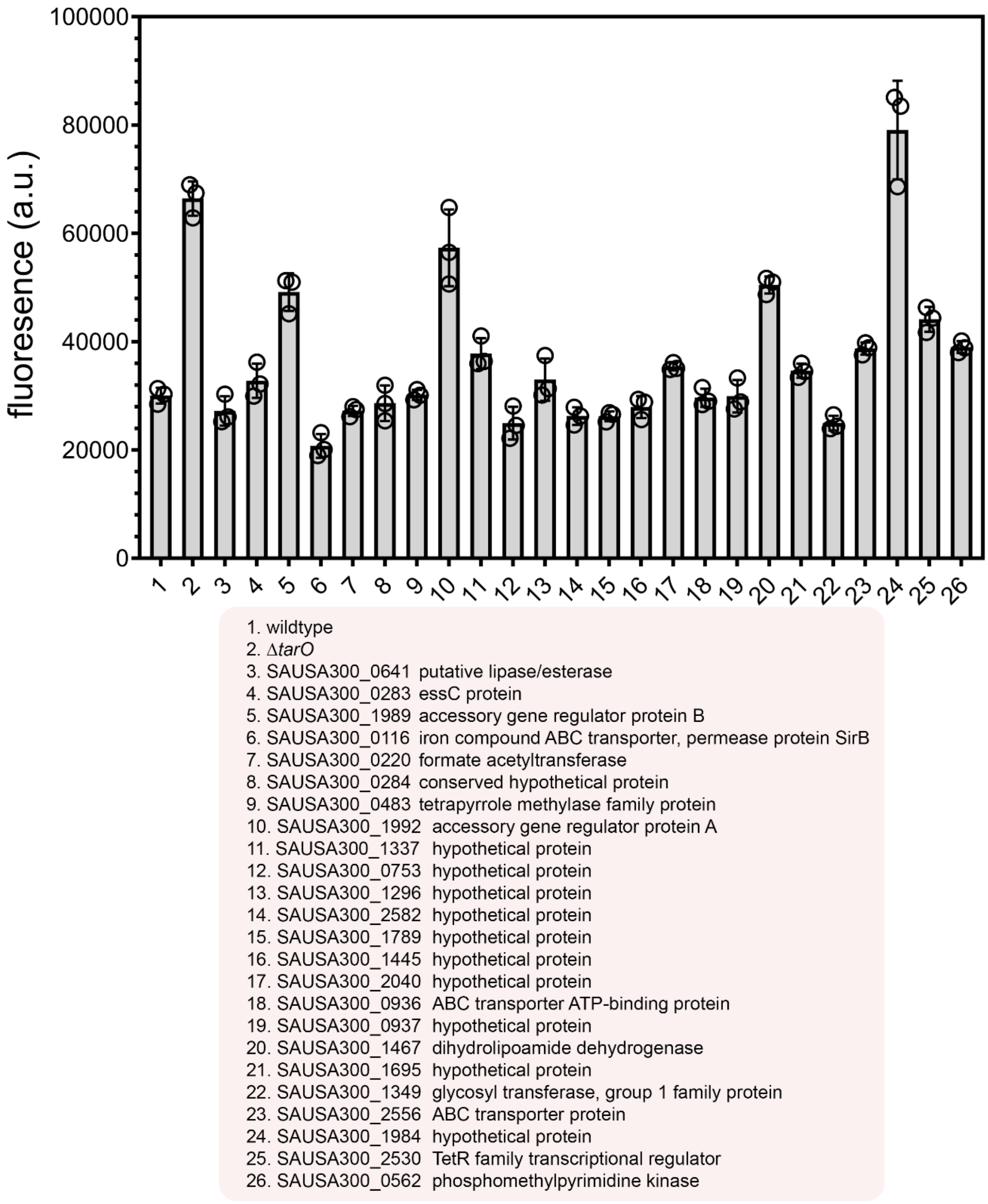


**Figure S3.** Flow cytometry analysis of designated *S. aureus* strains selected from transposon screen. Cells treated overnight with 1 mM of **D-LysAz**, followed by a treatment with 25 μM of **DBCO-Fl**. Data are represented as mean +/- SD (n = 3).

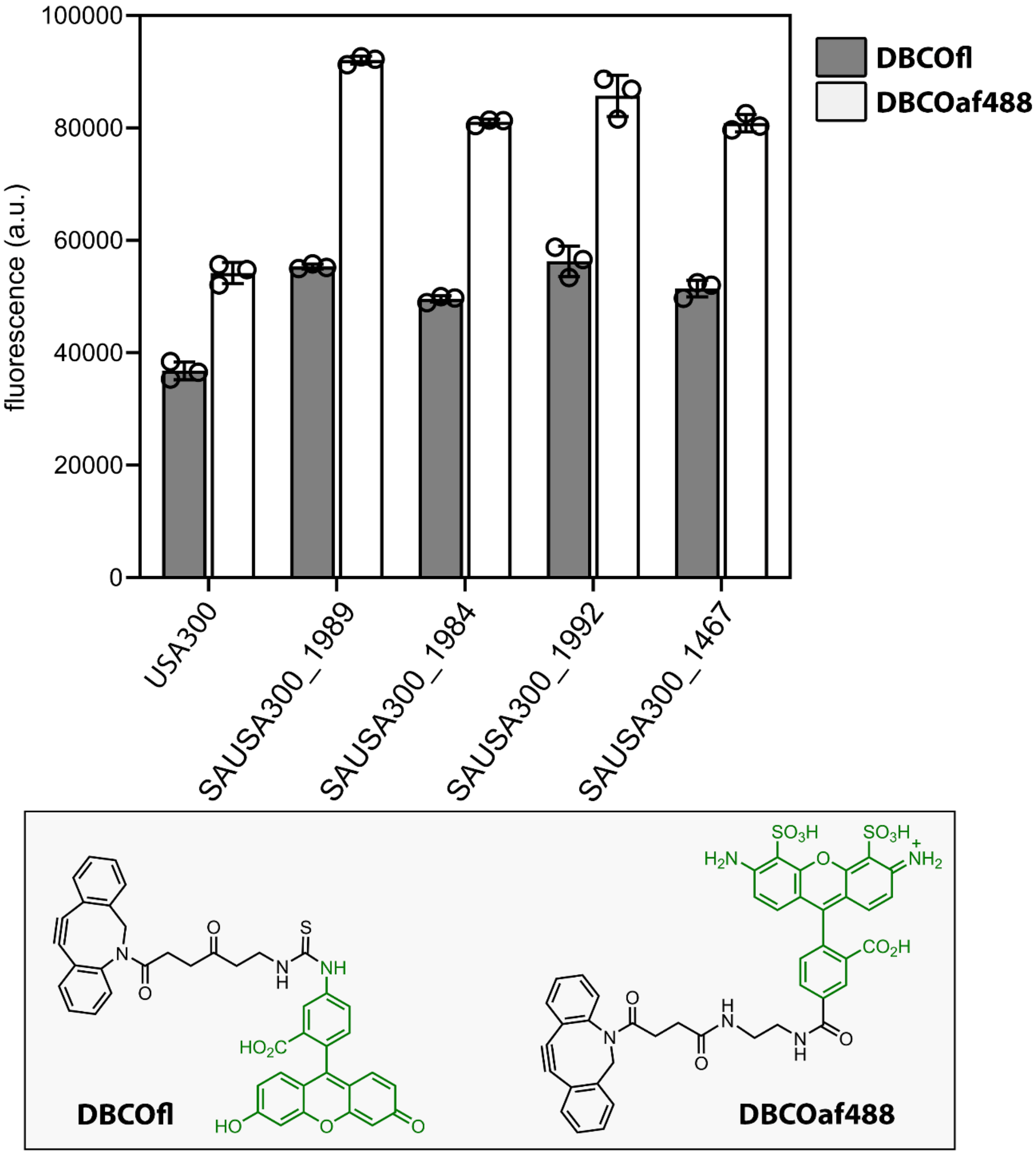
100000
80000
60000
40000
20000
0
fluorescence (a.u.)
DBCOfl
DBCOaf488
USA300
SAUSA300_1989
SAUSA300_1984
SAUSA300_1992
SAUSA300_1467
DBCOfl
DBCOaf488

**Figure S4.**

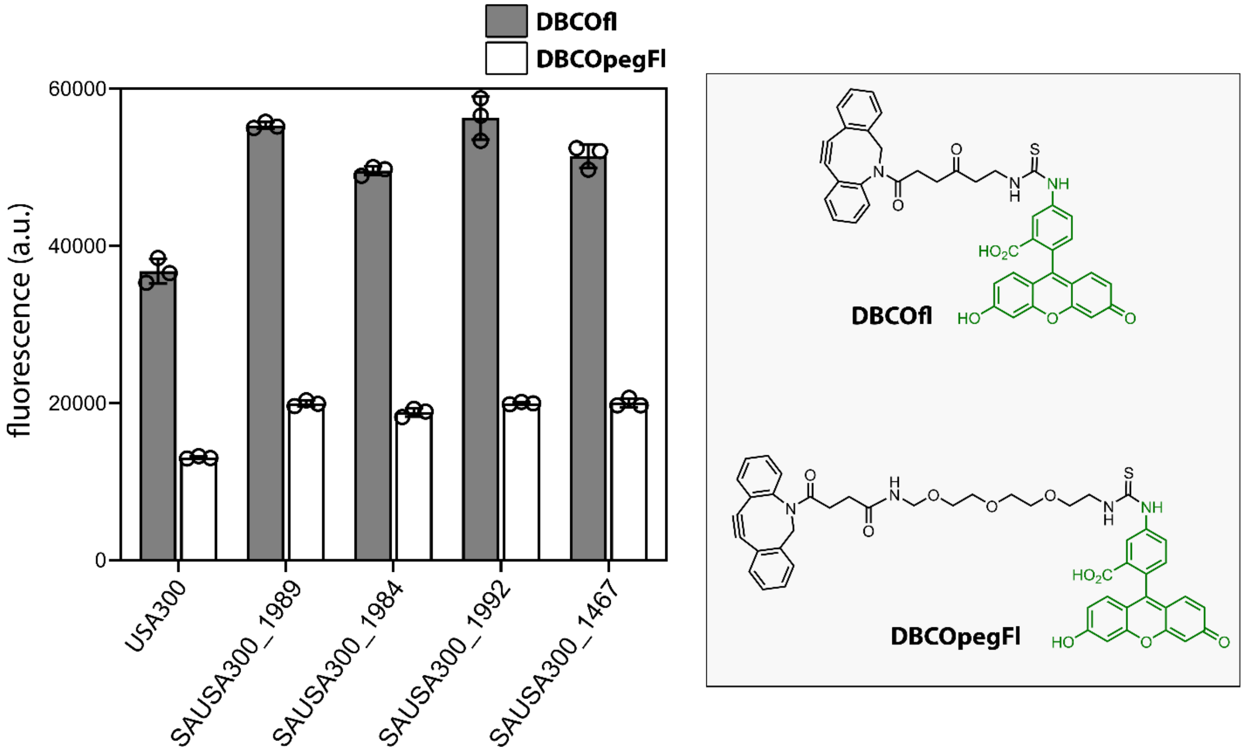

DBCOfl
DBCOpegFl
60000
40000
20000
0
fluorescence (a.u.)
USA300
SAUSA300_1989
SAUSA300_1984
SAUSA300_1992
SAUSA300_1467
DBCOfl
DBCOpegFl


**Figure S5.**

**Transposon Mutant Screen.** 35 μL of LB media was transferred by a liquid handling system into each well of the desired number of 384 well plates (120 μL max capacity). A 384 pin replicator was used to inoculate the prepared plates from the NTML glycerol stock plates. Inoculated plates were allowed incubate for 8 h with shaking at 37°C. 75 μL of fresh LB media containing 1mM of **D-LysAz** was transferred by a liquid handling system into each well of the desired number of 384 well plates (240 μL max capacity). After 8 h, the newly prepared plates were inoculated and allowed to incubate overnight with shaking at 37°C. The cells were then harvested at 4000 rpm and washed three times with 1X PBS. The pellets were then resuspended in the original culture volume (75 μL) and 10 μL of cells were transferred to plates containing **DBCO-Fl** for a final concentration of 25 μM and allowed to incubate for 30 minutes at 37°C. After 30 minutes the samples were fixed with 2% formaldehyde and analyzed using the Attune NxT flow cytometer as described above.

**Triplicate screen of hits**The triplicate screen of the identified hits was performed as described. LB media containing 1mM of **D-LysAz** was prepared. Identified hits grown overnight were added to the LB medium (1:100) and allowed to grow overnight at 37°C with shaking at 250 rpm. The cells were harvested at 4000 rpm and washed three times at the original culture volume with 1X PBS. The cells were then treated with 25 μM **DBCO-Fl** for 30 minutes at 37°C and protected from light. The samples were subsequently harvested at 4000 rpm and washed three times with 1X PBS followed by fixation with 2% formaldehyde in 1X PBS for 30 minutes. The cells were washed once more to remove formaldehyde and then analyzed using the Attune NxT flow cytometer as described above.

**Suseptibility of Transposon Mutants.** Identified hits from the triplicate screen in addition to *S. aureus* ATCC 25923 and USA300 were grown overnight. After growth cells were washed 3 times with 1X PBS and resuspended in buffer and the optical density at 600 nm was measured constituting the zero time point. Next 5 μg/mL of lysostaphin was introduced to each sample and allowed to incubate at 37°C with shaking at 250 rpm. The optical density at 600 nm was taken at each designated time point (5, 10, 30, 60, 120,180 mins). Percent initial optical density was calculated and graphed to represent the decrease in viability of cells over time.
